# Diet of Andean leaf-eared mice (*Phyllotis*) living at extreme elevations on Atacama volcanoes: insights from metagenomics, DNA metabarcoding, and stable isotopes

**DOI:** 10.1101/2024.07.23.604871

**Authors:** Claudio Quezada-Romegialli, Marcial Quiroga-Carmona, Guillermo D’Elía, Chris Harrod, Jay F. Storz

## Abstract

On the flanks of >6000 m Andean volcanoes that tower over the Atacama Desert, leaf-eared mice (*Phyllotis vaccarum*) live at extreme elevations that surpass known vegetation limits. What the mice eat in these barren, hyperarid environments has been the subject of much speculation. According to the arthropod fallout hypothesis, sustenance is provided by windblown insects that accumulate in snowdrifts (‘aolian deposits’). It is also possible that mice feed on saxicolous lichen or forms of cryptic vegetation that have yet to be discovered at such high elevations. We tested hypotheses about the diet of mice living at extreme elevations on Atacama volcanoes by combining metagenomic and DNA metabarcoding analyses of gut contents with stable-isotope analyses of mouse tissues. Genomic analyses of contents of the gastrointestinal tract of a live-captured mouse from the 6739 m summit of Volcán Llullaillaco revealed evidence for an opportunistic but purely herbivorous diet, including lichens. Although we found no evidence of animal DNA in gut contents of the summit mouse, stable isotope data indicate that mice native to elevations at or near vegetation limits (∼5100 m) include a larger fraction of animal prey in their diet than mice from lower elevations. Some plant species detected in the gut contents of the summit mouse are known to exist at lower elevations at the base of the volcano and in the surrounding Altiplano, suggesting that such plants may occur at higher elevations beneath the snowpack or in other cryptic microhabitats.

## 1. INTRODUCTION

Extreme high-elevation surveys of small mammals in the Central Andes have yielded live captures of numerous specimens of the Andean leaf-eared mouse *Phyllotis vaccarum* at elevations at or above the elevational limits of vegetation (>5,000 m)(Storz et al., 2024). One specimen was captured at 6,739 m (22,100 feet) above sea level on the very summit of Volcán Llullaillaco, a stratovolcano in the Central Andes that straddles the Argentina-Chile border (Storz et al., 2020). This summit specimen far surpasses previous elevational records for wild mammals in the Andes and Himalaya. Documentation of active burrows of *P. vaccarum* at >6,100 m on the flanks of Llullaillaco and the discovery of desiccated cadavers (‘mummies’) of *Phyllotis* on the summits of Llullaillaco and several neighboring >6,000 m volcanoes confirm that these mice inhabit extreme elevations well above the apparent limits of vascular plants (Halloy, 1991; Steppan et al., 2022; Storz et al., 2023, 2024). Evidence that high-elevation mice are living in an apparently barren world of rock, ice, and snow prompts numerous questions, perhaps none more basic than: What are they eating? In the perennial winter conditions that prevail at elevations >6,000 m, the scarcity of food poses a special physiological challenge for small endotherms like mice because of the energetic demands of thermoregulation. Moreover, leaf-eared mice in the genus *Phyllotis* do not hibernate, so the energetic challenge of sustaining endothermy in cold, hypoxic conditions is especially acute (Storz & Scott 2019).

It has been suggested that windblown arthropods and/or vegetation could provide a source of food for animals living at elevations that exceed the limits at which green plants grow (the ‘aolian zone’) (Swan, 1961, 1992). According to this hypothesis, the transport of airborne nutrients from lower elevations sustains life on the upper reaches of a volcano like Llullaillaco similar to the way that the fallout of organic detritus from upper layers of the water column sustains life in the aphotic zone of the deep ocean. Aolian deposits of windblown arthropods along the lee edge of mountain summits and ridgelines can attract birds and other insectivorous animals that normally forage at much lower elevations (Antor, 1995; Spalding, 1979). If aolian deposits of windblown arthropods (‘arthropod fallout’) help sustain populations of *P. vaccarum* at extreme elevations on Atacama volcanoes, we would expect arthropods to constitute a much larger part of their diet than at lower elevations where the species is mainly granivorous and frugivorous (Bozinovic and Rosenmann 1988; López-Cortés et al. 2007; Sassi et al. 2017). On the Quinghai-Tibetan Plateau, high elevation plateau pikas (*Ochotona curzoniae*) exploit the feces of yak (*Bos grunniens*) as a food source (Speakman et al., 2021). If *Phyllotis* mice practice a similar form of interspecific coprophagy, feces from Andean camelids such as vicuña (*Lama vicugna*) and guanaco (*Lama guanicoe*) would provide the most readily available source. Although neither vicuña nor guanaco typically spend much time above the elevational limit of vegetation, both species are known to traverse mountain passes at elevations >5,500 m in the Central Andes (J. Storz, personal observation). It is also possible that the mice feed on lichen that grows on rock substrates (saxicolous lichen) or some cryptic form of vegetation that is not currently known to occur at such extreme elevations.

Here we test the above-mentioned hypotheses by conducting metagenomic and metabarcoding analyses of gut contents from the world-record specimen of *P. vaccarum* that was live-captured on the summit of Llullaillaco at 6,739 m. Since the metagenomic approach involves high-throughput sequencing of all DNA extracted from a sample without PCR enrichment of specific markers, it is not biased by *a priori* expectations about which taxonomic groups to expect and is therefore well-suited to dietary assessments of omnivorous species (Chua et al., 2020). DNA metabarcoding complements the metagenomic approach and can be used to estimate the diversity and relative abundance of different items in the diet (Deagle et al., 2019; Stapleton et al., 2022).

To complement the metagenomic and metabarcoding analysis of the summit specimen, we conducted a stable isotope analysis of liver samples from a larger sample of wild-caught mice from a broad range of elevations in the Chilean Altiplano and Puna de Atacama (2,370-6,739 m). We used stable isotope values of three key elements (carbon, nitrogen and sulfur) from liver tissue to characterize the diet of *P. vaccarum* over a timespan of weeks to months. In the livers of small mammals, isotopic half-lives are <1 week for both carbon (δ^13^C) and nitrogen (δ^15^N) and we expect a similar half-life for sulfur (δ^34^S). Examination of stable isotope values permits inferences about several key components of trophic ecology.

Carbon stable isotopes (δ^13^C) reflect the relative consumption of food derived from different sources of primary production ( Cerling et al., 1997; DeNiro & Epstein, 1978). The stable isotope of nitrogen is typically used as an indicator of consumer trophic position (Vanderklift & Ponsard, 2003; Quezada-Romegialli et al., 2018) but can also be used to discriminate between consumption of food from distinct habitats (Harrod et al., 2005). For example, it should be possible to assess the extent to which mice rely on lichen at high elevations. Lichens are typically very ^15^N-depleted relative to terrestrial plants (Fogel et al., 2008; Lee, Lim & Yoon, 2009; Pinho et al., 2017), and this holds true for lichens from high elevations (Biazrov, 2012; Marris et al., 2019; Szpak et al., 2013) and volcanic fumeroles (Tozer *et al*. 2005). If lichen forms an important part of the diet of *P. vaccarum* at high elevations, we would expect to observe negative δ^15^N values. Sulfur stable isotopes (δ^34^S) are also useful indicators of consumer habitat use as values measured from plants and their consumers often exhibit high levels of spatial variation across biogeochemical gradients (Krouse et al., 1991; Nielsen et al., 1991), as might be expected along the flanks of an historically active volcano like Llullaillaco.

By combining metagenomics, metabarcoding, and stable-isotope analyses, we tested several hypotheses about the diet of mice living at extreme elevations. The arthropod fallout hypothesis would be supported by the presence of DNA from insects or other arthropods that could be blown upslope, and isotopic estimates of trophic position would be higher for mice living at especially high elevations on the flanks or summit of the volcano in comparison with those from lower elevations in the surrounding Altiplano. Interspecific coprophagy (or scavenging) would be supported by the presence of DNA from vicuña, guanaco, or other co-distributed mammals. Lichenivory would be supported by the presence of DNA from lichen-associated fungi or green algae and especially low ^15^N values in mice from high elevations. The particular plants detected in the gut contents of the summit mouse may suggest that some plant species actually occur at much higher elevations than currently assumed, but they may be sparsely distributed in cryptic microhabitats (e.g., in rock crevices or under the snowpack). If that is the case, then the diet of mice living at >6,000 m may include a subset of the same plants that mice feed on at lower elevations.

## 2. MATERIALS AND METHODS

### 2.1 Sampling

We live-captured all mice using Sherman live traps and other methods described in Storz et al., (2020, 2024). We collected mice from a broad range of elevations in the Altiplano/Puna de Atacama ecoregions, from 2,340 m in the Atacama Desert to the 6,739 m summit of Volcán Llullaillaco (Figure 1). We sacrificed mice in the field, prepared them as museum specimens, and preserved liver tissue in 95‰ ethanol as source material for the stable isotope analysis. For the *P. vaccarum* specimen captured on the summit of Volcán Llullaillaco, we preserved the entire gastrointestinal tract in ethanol as a source of DNA for metagenomic and metabarcoding analyses. All mouse specimens are housed in the Colección de Mamíferos of the Universidad Austral de Chile, Valdivia, Chile.

**Figure 1.**
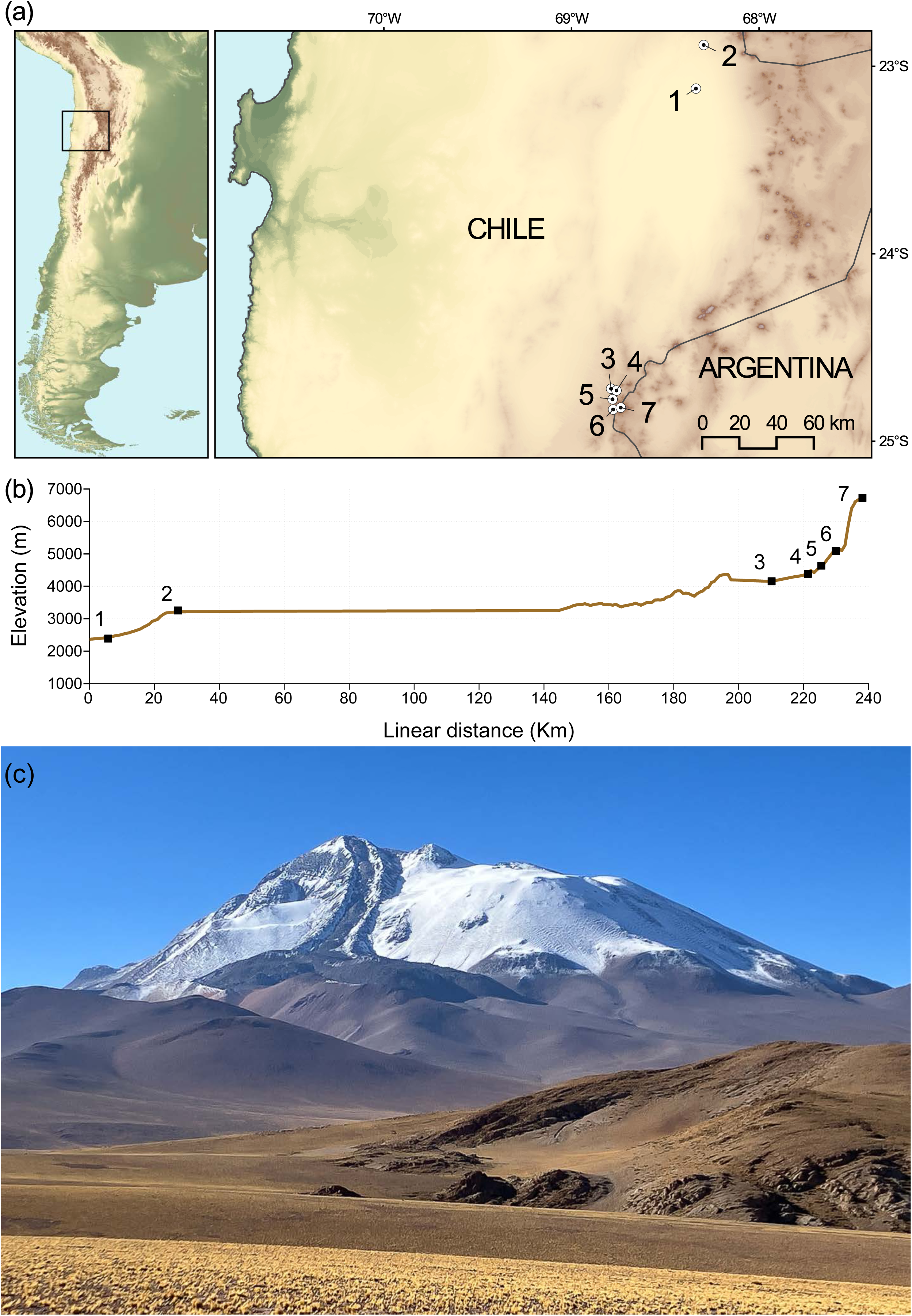
Sampling of *Phyllotis vaccarum* across an elevational gradient in the Altiplano and Puna de Atacama of northern Chile, Regiόn de Antofagasta. (a) Map of seven collection localities on the flanks of Volcán Lullaillaco and the surrounding Altiplano. (b) Elevational profile of sampling transect, with sampling localities 1 to 7 shown in ascending order of elevation, from 2370 m (site 1) to 6739 m (the summit of Volcán Lullaillaco, site 7). (c) Northwest face of Volcán Llullaillaco (24°43.21′S, 68°32.22′W). Photo was taken from a point ∼15 km northwest of the summit. Photo: J.F. Storz.

We collected all mice in accordance with permissions to JFS and GD from the following Chilean government agencies: Servicio Agrícola y Ganadero (SAG, Resolución exenta # 6633/2020), Corporación Nacional Forestal (CONAF, Autorización # 171219), and Dirección Nacional de Fronteras y Límites del Estado (DIFROL, Autorización de Expedición Científica #68/2020). We handled all mice in accordance with protocols approved by the Institutional Animal Care and Use Committee (IACUC) at the University of Nebraska (project ID: 1919).

### 2.2 Dissection of gastrointestinal tract

For the mouse captured on the summit of Volcán Llullaillaco (UACH8291), we extracted DNA from contents of the stomach for metagenomic sequencing and DNA metabarcoding. We also dissected the lower gastrointestinal tract into 13 adjoining sections, the cecum and 12 consecutive segments of the colon, ordered from the outlet of the cecum to the rectum (Figure 2), and we extracted DNA from the contents of each section for additional DNA metabarcoding analysis. This approach allowed us to examine temporal changes in the mouse’s diet, as determined by gut passage times: the stomach and cecum contain food items ingested within a few hours of its capture, while the colon sections and rectum potentially contain food ingested within the previous two or three days. The metagenomic sequencing represents an unbiased approach to characterize the stomach contents of the mouse while the metabarcoding analysis is designed to test specific hypotheses about the animal’s diet (arthropod fallout, interspecific coprophagy, lichenivory, or herbivory at elevations that surpass assumed vegetation limits).

**Figure 2.**
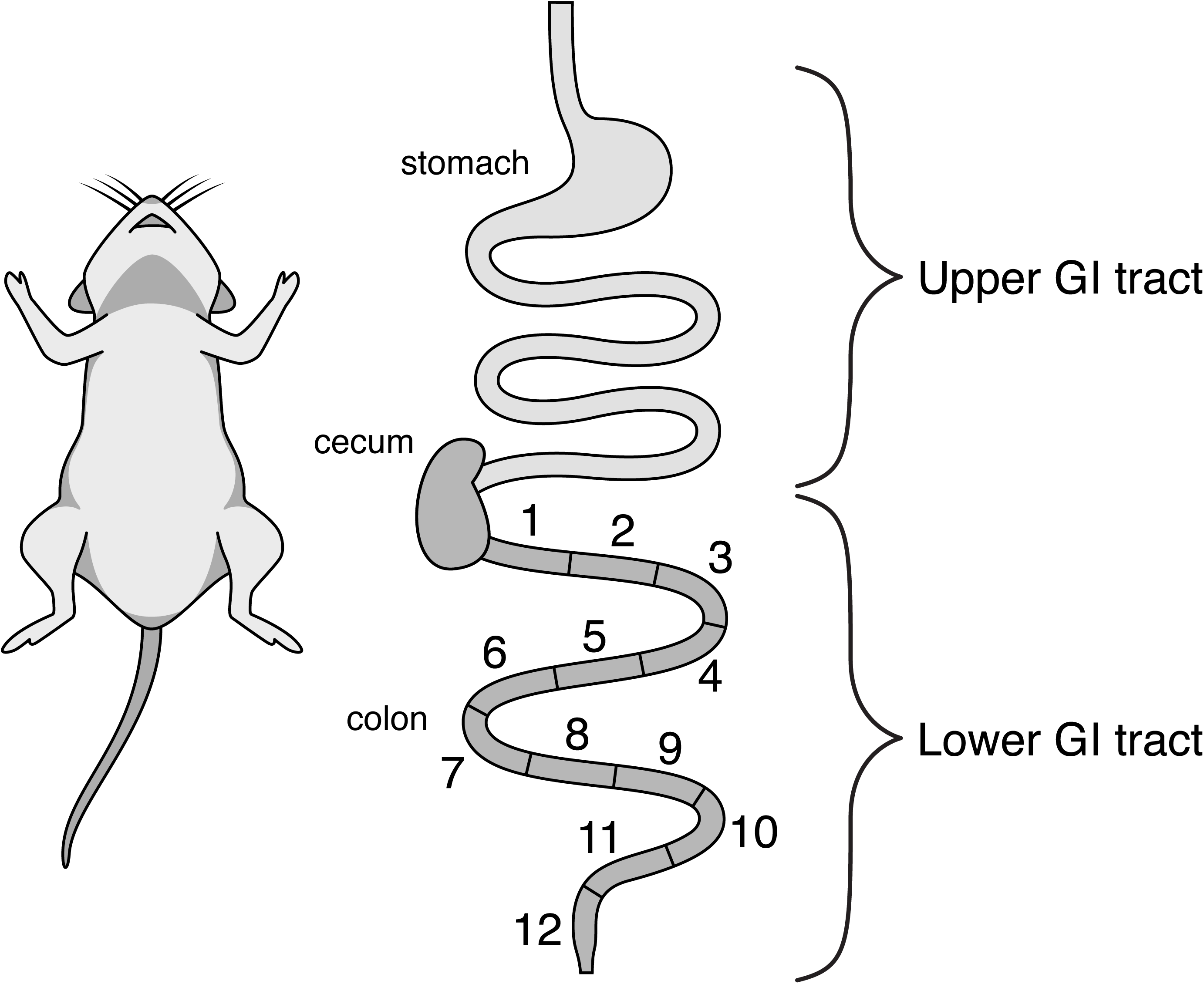
Schematic figure of the dissected portions of the gastrointestinal tract of the mouse from the summit of Volcán Llullaillaco. Contents of the upper digestive tract (stomach) were analyzed via metagenomics and DNA metabarcoding. Contents of the lower digestive tract (separated into the cecum and 12 consecutive segments from the outlet of the cecum to the rectum) were analyzed via DNA metabarcoding.

### 2.3 Metagenomic sequencing

We sent contents of the dissected sections of the gastrointestinal tract to Azenta Life Sciences (South Plainfield, NJ, USA) for metagenomic analysis. Genomic DNA was isolated using the NucleoMag DNA Microbiome Kit (Takara Bio, Shiga, Japan) and was quantified using a Qubit

2.0 Fluorometer (ThermoFisher Scientific, Waltham, MA, US). NEBNext® Ultra™ II DNA Library Prep Kit for Illumina (New England Biolabs, Ipswich, MA, USA) was used for library preparation following manufacturer’s recommendations. Briefly, genomic DNA was fragmented by acoustic shearing with a Covaris S220 instrument, followed by end-repair. Adapters were ligated after adenylation of the 3’ends followed by enrichment by a limited cycle PCR. DNA libraries were quantified using Qubit 2.0 Fluorometer and by real time PCR (Applied Biosystems, Carlsbad, CA, USA). Sequencing libraries were sequenced on an Illumina HiSeq instrument using a 2 x 150bp Paired End (PE) configuration. Image analysis and base calling were conducted using the HiSeq Control Software (HCS). Raw BCL files were converted to FastQ files and de-multiplexed using bcl2fastq v.2.1.9 (Illumina), keeping only >Q30 reads with 150 bp in length. A *de novo* approach was followed for assembling reads using Spades v3.10 (Bankevich et al., 2012), with a minimum contig length of 1,000 bp and using the newly assembled genome of *Phyllotis vaccarum* (Storz et al., 2023) as a reference genome to discard reads aligned to the host. QUAST (Gurevich et al., 2013) was used to generate statistics and EMBOSS tools getorf was used to find the open reading frames within the *de novo* assembled genome. BLAST+ (v.2.6.0) (Altschul et al., 1990) was used to query assembled contigs in the nucleotide database of Genbank.

### 2.4 DNA metabarcoding analysis and primer selection

The stomach, cecum, and 12 consecutive segments of the colon were sent to MrDNA (www.mrdnalab.com) for DNA extraction and metabarcoding analysis. We identified and discarded false positives using extraction blanks and multiple PCR replicates to avoid noise from spurious amplification (Taberlet et al., 2018, Table 1). We used the following primer pairs for specific taxonomic groups (Table 1): for plants, we amplified (i) the P6 loop of the chloroplast *trnL* (UAA) intron using primers P6-trnLF: 5’-GGG CAA TCC TGA GCC AA-3’ and p6-trnLR: 5’-CCA TTG AGT CTC TGC ACC TAT C-3’ (Taberlet et al 2007), and (ii) the internal transcribed spacer 2 (ITS2) of nuclear ribosomal DNA using primers S2F: 5’-ATGCGATACTTGGTGTGAAT-3’ and S2R: 5’-GACGCTTCTCCAGACTACAAT-3’ (Chen et al 2010); for eukaryotic algae and cyanobacteria, we amplified domain V of the 23S plastid rRNA gene using primers p23SrV_f1 5’-GGA CAG AAA GAC CCT ATG AA-3’ and p23SrV_r1 5’-TCA GCC TGT TAT CCC TAG AG-3’ (Sherwood & Presting, 2007); for fungi (Ascomycota and Basidiomycota), we amplified the internal transcribed spacer (ITS) of nuclear ribosomal DNA using primers: ITS1-F 5’-CTT GGT CAT TTA GAG GAA GTA A-3’ (Gardes & Bruns, 1993) and 5’-GCT GCG TTC TTC ATC GAT GC-3’ (White et al 1990); for metazoans, we amplified the cytochrome c oxidase subunit I using primers: mlCOIintF: 5’-GGW ACW GGW TGA ACW GTW TAY CCY CC-3’ (Leray et al., 2013) and jgHCO2198: 5’-TAI ACY TCI GGR TGI CCR AAR AAY CA-3’ (Geller et al., 2013); and for invertebrates, we amplified the cytochrome c oxidase subunit I using primers: fwhF2: 5’-GGD ACW GGW TGA ACW GTW TAY CCH CC-3’ (Vamos et al., 2017) and EPTDr2n: 5’-CAA ACA AAT ARD GGT ATT CGD TY-3’ (Leese et al., 2021).

**Table 1.** Summary statistics for metabarcoding sequence reads for each of the dissected portions of the gastrointestinal tract of the *Phyllotis vaccarum* specimen captured on the summit of Volcán Llullaillaco (see Figure 2).

| Sample | Input reads | Filtered | Denoised forward | Denoised reverse | Merged | Non-chimeric |
| --- | --- | --- | --- | --- | --- | --- |
| Extraction blanks | 2.009 | 1.173 | 1.150 | 1.142 | 13 | 13 |
| Stomach | 755.758 | 350.866 | 349.765 | 349.840 | 242.553 | 237.567 |
| Cecum | 742.138 | 321.525 | 320.278 | 319.642 | 229.464 | 226.427 |
| C1 | 548.838 | 283.847 | 283.023 | 282.946 | 213.947 | 204.749 |
| C2 | 745.252 | 370.547 | 369.633 | 369.303 | 271.283 | 261.741 |
| C3 | 634.524 | 324.615 | 323.596 | 323.317 | 226.881 | 224.844 |
| C4 | 551.463 | 254.049 | 253.346 | 253.147 | 206.205 | 203.786 |
| C5 | 559.460 | 272.415 | 271.609 | 271.402 | 159.567 | 158.241 |
| C6 | 624.732 | 301.630 | 300.660 | 300.059 | 193.624 | 188.859 |
| C7 | 670.502 | 313.308 | 311.935 | 311.397 | 214.056 | 205.772 |
| C8 | 760.648 | 397.214 | 395.953 | 395.543 | 287.349 | 282.642 |
| C9 | 731.646 | 367.289 | 365.790 | 365.438 | 275.122 | 269.431 |
| C10 | 658.965 | 318.646 | 317.186 | 316.856 | 215.254 | 212.705 |
| C11 | 702.488 | 340.687 | 339.058 | 338.399 | 224.913 | 219.703 |
| C12 | 798.662 | 390.642 | 388.897 | 388.447 | 304.740 | 300.431 |
| **TOTAL** | **9.487.085** | **4.608.453** | **4.591.879** | **4.586.878** | **3.264.971** | **3.196.911** |

### 2.5 Bioinformatic processing – metabarcoding

Reads were analyzed separately for each primer in R v 4.3.1 (R Core Team, 2023) using RStudio 2023.12.1 (RStudio Team, 2023) in dada2 (Callahan *et al*. 2016), after filtering (maxEE = 2, Q-scores >30; Edgar & Flyvbjerg, 2015) and discarding reads with Ns. Error model calculation, read correction, and read merging and removal of chimeric sequences was performed using default settings. Amplicon sequence variants (ASVs) identified by dada2 were assigned to taxa of origin using BLAST+ (v.2.6.0) (Altschul et al., 1990) and the GenBank database.

### 2.6 Stable isotope analysis

Ethanol-preserved liver tissues were rinsed in distilled water and freeze-dried for 48 h. Once freeze-dried, samples were ground to a fine powder using a laboratory bead-beater. Ground samples were weighed in 8 x 5 mm pressed standard weight tin capsules using a high precision microbalance (repeatability = 0.0008 mg). Elemental percentages of carbon, nitrogen, sulfur, and stable isotope ratios (δ^13^C, δ^15^N and δ^34^S) were measured using a Pyrocube elemental analyser (Elementar, Langenselbold, Germany) linked to a visION continuous-flow isotope ratio mass spectrometer (Elementar, Langenselbold, Germany) at the Universidad de Antofagasta Stable Isotope Facility (UASIF), Chile. Stable isotope ratios are expressed using δ notation and are reported in units of per mil (‰) relative to the following standards: Vienna Pee Dee Belemnite for carbon, air for nitrogen, and Vienna Canyon Diablo Troilite for sulfur. International standards were used in each batch to provide a multi-point calibration using the ionOS software package v4.1.005 (Elementar, Langenselbold, Germany). Certified reference material USGS40 and USGS41a were used for carbon and nitrogen and IAEA-SO-5, IAEA-SO-6 and IAEA-S2 for sulfur. Repeated analysis of standards showed analytical errors (± 1 SD) of ± 0.04 ‰ for δ^13^C, ± 0.06 ‰ for δ^15^N, and ± 0.6 ‰ for δ^34^S. We used two calibration standards, a) sulfonamide (Elementar, Germany) and b) an in-house standard (rainbow trout dorsal muscle) to correct for instrument drift.

### 2.7 Statistical analyses – stable isotopes

Liver is commonly used as a lipid-storage organ in vertebrates and liver lipid content can vary significantly among individuals according to variation in nutritional state and physiological condition. Lipids formed through *de novo* biosynthesis are isotopically lighter in δ^13^C values compared to proteins and the dietary sources from which they were formed (DeNiro & Epstein, 1977). If the isotopic effect of these lipids is not accounted for, they can affect assessment of consumer δ^13^C values. Furthermore, as the livers analyzed here were preserved in ethanol, they may have undergone some partial uncontrolled lipid extraction prior to analysis. Variation in individual lipid content can affect comparisons of δ^13^C values and it is therefore common to use chemical treatments to remove lipids prior to stable isotope analyses. However, the chemical treatment can affect estimated values of other stable isotopes from the same sample. Another possible solution is to use an arithmetic correction that relies on a predictable relationship between lipid content and the elemental ratio between carbon and nitrogen (C:N) in the sample (Kiljunen et al., 2006; Logan et al., 2008). Javornik et al. (2019) found small effects of ethanol storage on δ^13^C values in mammalian liver but reported no preservation effects on δ^15^N or δ^34^S values. Javornik et al. (2019) reported that C:N values decreased after ethanol storage but suggested that lipid-free ^13^C values could be reliably estimated mathematically from the C:N ratio. In our samples, liver C:N ratios varied considerably (range = 3.2-5.7, mean ± SD = 3.8 ± 0.5, *n* = 41). Liver C:N ratios were lower in mice captured at higher elevations (*r* = -0.36, *n* = 40, *P* = 0.024). As there was also a negative relationship between C:N and δ^13^C within samples from the same collection locality, we estimated lipid-corrected δ^13^C values using Equation 1a from Logan et al. (2008), resulting in a mean (± SD) isotopic shift of 1.1 ± 0.6‰. All δ^13^C data that we report are lipid-corrected, but liver δ^15^N and δ^34^S data are shown without correction.

Collection sites spanned ∼4400 m of elevation and therefore exhibited considerable variation in vegetation cover and plant species composition. We therefore examined how stable isotopes varied within the dataset by plotting each of the stable isotopes against elevation. We then used PERMANOVA (non-parametric permutation-based equivalent of ANOVA) to examine whether stable isotope values varied among capture sites. Since we collected a single individual from site 7 (the summit of Llullaillaco), it was not included in these comparisons. Although PERMANOVA is typically used for multivariate comparisons (MANOVA), it can also be used to make robust univariate comparisons. Finally, to assess the ability of the stable isotope analysis to assign mice to capture location, and to identify mismatches that may be indicative of recent dispersal, we used canonical analysis of principal coordinates (CAP), a distance-based equivalent of discriminant function analysis (Anderson & Willis, 2003). This approach uses multivariate data (e.g., δ^13^C, δ^15^N and δ^34^S values) to discriminate between groups defined by elevation of capture sites. This approach also allowed us to infer the possible origin of the summit mouse from Volcán Llullaillaco (site 7). We grouped mice in bins based on their capture elevation (2000-3000 m, 3000-4000 m, 4000-5000 m, and >5000 m) and we used δ^13^C, δ^15^N and δ^34^S as dependent variables. We used a leave-one-out classification approach to examine relative classification success, and we then used the model to identify the elevational range that provided the best match to values from the Llullaillaco summit mouse. The ability of the CAP model to statistically discriminate between groups was estimated via permutation (*n* = 9999). PERMANOVA and CAP were both run in the PERMANOVA+1 add on to PRIMER 7 (Anderson et al., 2008; Clarke & Gorley, 2015).

We estimated the trophic position of *P. vaccarum* at each site using liver δ^15^N values with those of primary producers collected across a similar (but truncated) elevational range (Díaz et al., 2016). This approach (Cabana and Rasmussen 1996) allows the indirect calculation of consumer trophic position (TP): TP = λ + (δ^15^N_Consumer_ − δ^15^N_Baseline_)/TDF, where λ is the trophic position of the baseline taxon, δ^15^N_Consumer_ is the nitrogen isotopic value of mice at a given site, δ^15^N_Baseline_ is the nitrogen isotopic value of the baseline at that site, and TDF is the mean ± SD nitrogen trophic enrichment factor (TDF) for mouse liver (here we use 4.3 ± 0.2 ‰ from Arneson and MacAvoy (2005)). We used plants as our baseline (λ =1) based on data from Díaz et al. (2016), which were collected in the same region as our study, over an elevational range of 2670-4480 m. Plant δ^13^C and δ^15^N values exhibited considerable variation across elevations (Figure 6), and we placed plants into broad elevational intervals (2000-3000 m, 3000-4000 m, 4000-5000 m, and >5000 m). We then used values from the closest elevational interval to estimate mouse trophic position at each capture site using *tRophicPostion* 0.8.0 (Quezada-Romegialli et al., 2018) in R 4.2.3 (López-Cortés et al., 2007; R Core Team, 2023). Briefly, *tRophicPosition* uses a Bayesian approach to estimate trophic position for a population of consumers while accounting for variation in consumer and baseline isotope values. For most sites we use the *onebaseline* model (assuming a single baseline) but we used the *twoBaselines* model for mice from site 2. This is because the plants from the 3000 – 4000 m interval showed a bimodal distribution of δ^13^C values, which indicates the presence of plants using different photosynthetic pathways (e.g. C3, C4/CAM). Since these groups also showed evidence for a non-normal distribution of δ^15^N values, we used the twoBaselines full model, which also uses baseline δ^13^C. For all model runs we used the following parameters: chains = 3, number of adaptive iterations = 1 000, iterations = 20 000, burn-in = 1 000, thinning = 10. In case of the summit mouse from Volcán Llullaillaco we developed an individual model to calculate trophic position, as *tRophicPosition* v 0.8.0 currently provides only population-level estimates of TP. This new model with a one baseline approach was implemented in *greta* (Golding, 2019) which allows the calculation of TP at the individual level. We modelled the baseline for the summit mouse as having a mean and standard deviation of δ^15^N values of plants >5,000 m with a normal distribution for the mean and a Cauchy distribution for the SD, with a location of plants δ^15^N SD a scale of 3 and truncated from 0 to infinite. In this analysis λ is 1, the TDF was modelled as having a normal distribution with a mean of 4.3 and SD of 0.2 (Arneson & MacAvoy, 2005). We calculated 10 000 samples, with a thinning of 10, 1 000 samples as warmup and 16 chains.

Due to the selective retention of heavier isotopes during the assimilation of food, consumers are typically isotopically ‘heavier’ than their food (DeNiro & Epstein, 1978; DeNiro & Epstein, 1981). These diet-tissue shifts are referred to as trophic enrichment or trophic discriminations, and are typically estimated in experimental settings. Arneson and MacAvoy (2005) provided empirical estimates for trophic discrimination factors (TDFs) in liver from groups of laboratory mice fed diets that differed in the origin of their protein and carbohydrate components. In their control diet, where carbohydrates and proteins originated from the same source, mean ± SD TDFs were 0.7 ± 0.3 ‰ for carbon (Δ^13^C), 4.3 ± 0.2 ‰ for nitrogen (Δ^15^N), and -2.1 ± 0.1 ‰ for sulfur (Δ^34^S). As such, we expect mouse livers to have δ^13^C and δ^15^N values that are ∼1% and ∼4 % higher, respectively, than their long-term average, in combination with δ^34^S values ∼2 % lower than the long-term average.

## 3. RESULTS AND DISCUSSION

### 3.1 Metagenomics

For the stomach DNA sample of the Llullaillaco summit mouse, we sequenced a total of 423,477,275 reads, yielding 127,043 Mbases, with 92.48% of reads ≥ q30 and a mean quality score of 35.69. We assembled a total of 9,138 contigs ≥1,000 bp in length (21,188,348 bp), with a maximum length of 103,345 bp. Out of 991 contigs that were identified at the order level and above, the vast majority were assigned to super kingdom Bacteria (699 contigs), with Proteobacteria (563 contigs) and Firmicutes (99) as the dominant phyla. Only 1.3% of contigs (*n*=13) were assigned to plants (clade Streptophyta, class Magnoliopsida [= dicotyledons]), and all were assigned to a single representative of the coca family, Erythroxylaceae (*Erythroxylum novagranatense*). This shrub species is widely cultivated in South America because its leaves are a rich source of the psychoactive alkaloid, cocaine. In the stomach contents of the summit mouse, we detected no traces of DNA from arthropods nor from vicuña, guanaco, or other potentially co-distributed Andean mammals.

### 3.2 Metabarcoding

We sequenced a total of 9,487,085 reads as part of the DNA metabarcoding analysis, maintaining 3,196,911 non-chimeric reads after filtering, denoising, and merging reads for all primer combinations and all samples (Table 1, Tables S1-S6). The P6 loop of the chloroplast *trnL* (UAA) intron marker (Figure 3a,b) yielded an average of 56,507 ± 22,886 (SD) non-chimeric processed reads for each of the dissected sections of the gastrointestinal tract. Consistent with results of the metagenomic analysis, sequences assigned to the family Erythroxylaceae (*Erythroxylum novagranatense*) predominated in samples from each section, from the stomach to the C12 portion of the colon (Figure 3a). Sequences of *E. novagranatense* represent 97.3% of the 886,078 sequences derived from all surveyed sections of the gastrointestinal tract. In addition to representatives of Erythroxylaceae, we detected representatives of Amaryllidaceae (0.4% of the total), Poaceae (0.3%), Pinaceae (0.3%), and Malvaceae (0.1%), and Fabaceae and Juglandaceae, which together accounted for 0.05% of the total (Figure 3b).

**Figure 3.**
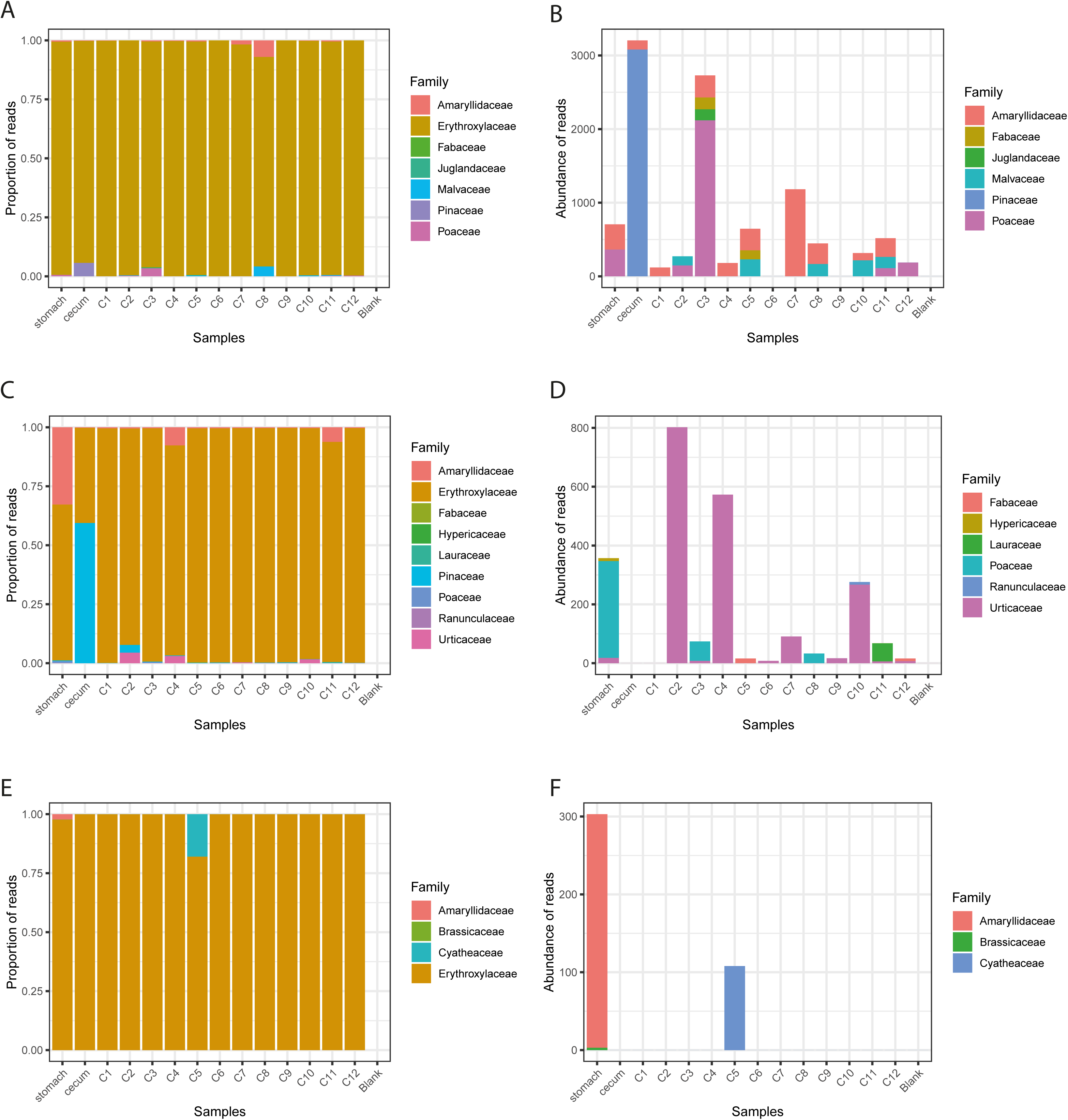
Taxonomic composition of reads identified through metabarcoding for the stomach, cecum and 12 consecutive portions of the lower GI from the outlet of the cecum to the rectum. (A) Proportion of sequence reads per taxon for the marker *trn* and (B) proportion of total reads for the same marker with Erythroxylaceae excluded. (C) Proportion of sequence reads per taxon for the marker *ITS2* and (D) proportion of total reads for the same marker with Erythroxylaceae excluded. (E) Proportion of sequence reads per taxon for the marker *23S* and (F) proportion of total reads for the same marker with Erythroxylaceae excluded.

The internal transcribed spacer 2 (ITS2) marker yielded an average of 19,952 ± 7,762 non-chimeric reads for each section of the gastrointestinal tract, and Erythroxylaceae accounted for 87.2% of the 279,336 total reads. Sequences from Pinaceae (6.1% of the total), Amaryllidaceae (5.9%), Urticaceae (0.64%), Poaceae (0.15%) were also detected, along with traces of Laureaceae, Fabaceae, Ranunculaceae and Hypericaceae, which together accounted for 0.04% of total reads (Figure 3d).

The third marker that provided information for Streptophyta was domain V of the 23S plastid rRNA gene which yielded 1,596 ± 671 reads on average for all sections of the gastrointestinal tract. Again, Erythroxylaceae predominated, accounting for 98.2% of 22,352 total reads (Figure 3e). In addition to Erythroxylaceae and Amaryllidaceae (1.5% of total reads), the next two most abundant taxa were Cyatheaceae and Brassicaceae, which together accounted for <0.6% of total reads (Figure 3f).

For Fungi, the internal transcribed spacer (ITS) of the nuclear ribosomal DNA marker detected a wide variety of Ascomycota and Basidiomycota orders and families (Figure 4) in the 414,990 read total, with an average of 29,642 ± 12,846 reads for each section of the gastrointestinal tract. For Ascomycota, the most abundant families were Cladosporiacea (28.4% of the total reads for this Phylum), Pleosporaceae (20.2% of the total), Sacharomycetaceae (16.6% of the total) and Nectriaceae (14.1% of the total), while the remaining 24 families represent 20.6% of the total, with Phaeococcomycetaceae and Parmeliaceae (lichen-associated families) accounting for 1.6% of the total. For the Phylum Basidiocomycota, the orders Agaricales (40.7% of total reads for this Phylum) and Polyporales (36.5% of the total) represent the most abundant groups, whereas Psathyrellaceae (18.8% of the total), Polyporaceae (13.6% of the total), Agaricaceae (12.0% of the total), and the families Meripilaceae, Fomitopsidaceae and Hyphodermataceae (together accounting for 16.7% of the total) were the most abundant groups across all sections of the gastrointestinal tract.

**Figure 4.**
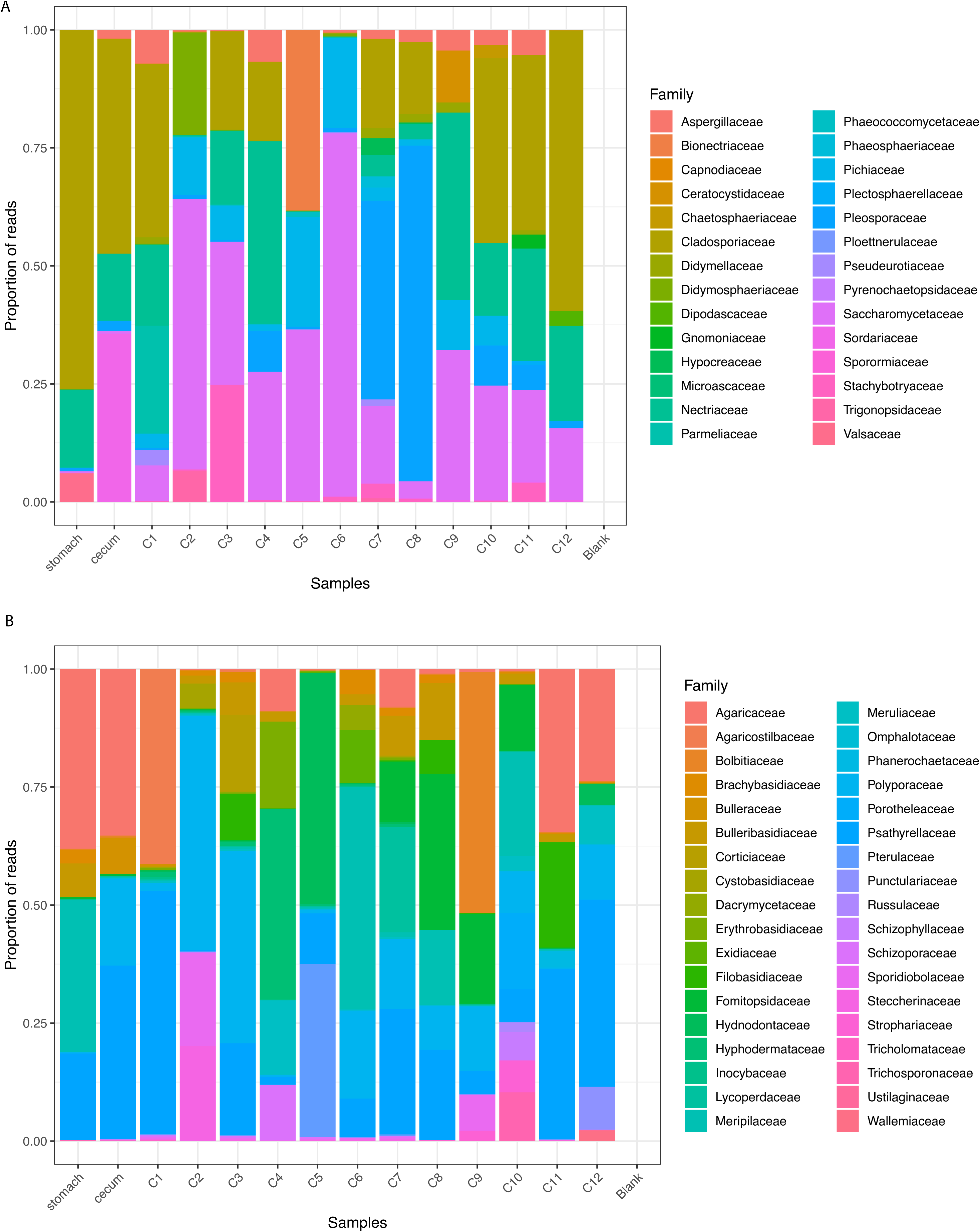
Taxonomic composition of Ascomycota (A) and Basidiomycota (B) at the Order and Family levels.

For Metazoans the marker cytochrome c oxidase subunit I identified 41 amplicon sequence variants (ASV), all of which were assigned to *Phyllotis*; none were derived from vicuña or guanaco. Nine ASV remain uncategorized. Finally, the marker cytochrome c oxidase subunit I specifically developed for arthropods did not detect any ASV for the group.

### 3.3 Diet of the summit mouse

One of the most puzzling results of the metagenomic and metabarcoding analyses is the predominance of Erythroxylaceae (the coca family) and Amaryllidaceae (the garlic family) in the stomach, cecum, and all 12 of the independently analyzed sections of the colon (Figure 3). Two representatives of Erythroxylaceae, *Erythroxylum argentinum* and *E. cuneifolia*, exist at elevations below ∼2000 m far to the east of Llullaillaco, but they do not occur in Andean desert or dry puna habitats. Coca was widely used throughout the Incan empire and is still used in indigenous Quechua and Aymara communities and, occasionally, by mountain climbers. Upon summiting a particular peak, there is a custom (especially among Argentine climbers) of leaving offerings to Pachamama, an Andean “Earth mother”, Gaia-type deity. Such offerings are left at the base of rock piles called ‘apacheta’ that serve as summit markers. A typical offering to Pachamama is a sprinkling of coca leaves or a small bag of such leaves at the base of the apacheta. This custom provides a ready explanation for the predominance of Erythroxylaceae in the gut contents of the summit mouse, which had presumably encountered just such an offering on the summit of Llullaillaco. The presence of garlic in the stomach contents of the summit mouse has a similar explanation. In the Argentine province of Salta (where the western portion of Llullaillaco is located), garlic is a traditional folk remedy for altitude sickness. Argentine climbers are known to chew cloves of garlic during their ascent. As is the case with any unchewed coca that climbers possess upon reaching the summit, it is also customary to leave leftover cloves of garlic at the base of the summit apacheta. The predominance of both coca and garlic in the gut contents of the Llullaillaco summit mouse suggest that climbers’ offerings to Pachamama on the summits of high Andean summits may sometimes serve as unintentional offerings to opportunistic *Phyllotis* mice living in an extremely food-scarce environment.

Aside from Erthroxylaceae and Amaryllidaceae, we also detected DNA representative of several plant families such as Fabaceae, Malvaceae, and Poaceae, that occur at high elevations at the base of Volcán Llullaillaco and in the surrounding Altiplano (Arroyo et al., 1988, 1998; Luebert & Gajardo, 1999; Marticorena et al., 2004). Within Fabaceae, the herb *Astragalus pusillus* was documented at elevations up to 4300 m on the flanks of Llullaillaco (Marticorena et al., 2004). Within Malvaceae, several perennial herbs such as *Cristaria andicola*, *Nototriche auricoma*, and *N. clandestina* occur at elevations between 4000-4500 m (Arroyo et al. 1998) and *Cristaria andicola* was documented as the most abundant plant species in the diet of *Phyllotis* at another altiplano study site in northern Chile (López-Cortés et al., 2007). Within Poaceae (=Gramineae), bunch grasses in the genus *Calamagrostis* (recognized as *Deyeuxia* in Arroyo et al. [1998] and Luebert & Gajardo [1999]) occur above 4000 m (Marticorena et al., 2004) and are also known to be included in the diet of *Phyllotis* from the Chilean Altiplano (López-Cortés et al. 2007). Although all of these plants seem plausible as potential sources of food for the summit mouse, there are no records of vascular plants or other vegetation above ∼5000 m on the flanks of Llullaillaco (Arroyo et al., 1988, 1998; Luebert & Gajardo, 1999; Marticorena et al., 2004; Storz et al., 2024; Vimercati et al., 2019), although it is also true that botanical surveys typically do not venture above such elevations, so we should be careful about interpreting absence of evidence as evidence of absence.

Given that sequences representative of Fabaceae, Malvaceae, and Poaceae were detected in the gut contents of the mouse captured at 6739 m, far above the apparent elevational limits of those plant taxa, there are three possible explanations to consider: (1) plant material is carried upslope by the wind and accumulates in sufficient quantities on the lee edge of ridge lines and snowdrifts to provide a source of sustenance for high-elevation mice (‘Aolian deposits’; Antor, 1995; Spalding, 1979; Swan 1961, 1992); (2) the plants in question actually occur at much higher elevations than previously thought (though they may be scarce and cryptic); or (3) the mouse was not a full-time summit resident, but rather a transient sojourner that had simply consumed the plant material at or near the base of the mountain some days prior to its capture. The former two hypothesis cannot be rejected, since few systematic plant surveys have been performed on Llullaillaco or other >6000 m volcanoes (Storz et al., 2024). In assessing the plausibility of the third hypothesis, it is important to note that the ∼1.6 km elevational distance between the summit of Llullaillaco (6739 m) and the apparent vegetational limit (∼5000-5100 m; Storz et al., 2024) translates into a linear distance of ∼5 km from any side of the volcano. Summiting the volcano from the vegetation limit is roughly equivalent to a direct-from-basecamp ascent, a feat that only the most elite mountain climbers could accomplish in a single day. We cannot rule out the possibility that mice undergo upslope/downslope dispersal on a seasonal basis, but such movements could certainly not occur on a daily basis. Moreover, in addition to the live capture of the *P. vaccarum* specimen on the Llullailllaco summit at 6739 m, video records and identification of active burrows of *P. vaccarum* between 6145 – 6205 m on the same volcano, and the discovery of desiccated cadavers and skeletal remains of numerous *P. vaccarum* on the summits of four different >6000 m volcanoes in the same mountain chain (Halloy, 1991; Steppan et al., 2023; Storz et al., 2020, 2023, 2024) provide a consilience of evidence suggesting that these extreme high-elevation mice are representative of resident populations. We think it is more likely that potential food plants exist at higher elevations, although they must be scarce and patchily distributed. The plausibility of this hypothesis is supported by the surprising discovery of bryophytes growing in association with active volcanic fumaroles near the summit of Volcán Socompa (Halloy, 1991), a 6051 m volcano located 47 km northeast of Llullaillaco along the Argentina-Chile border.

Primer set ITS detected sequences from families of two lichen-associated fungi, Phaeococcomycetaceae y Parmeliaceae (phylum Ascomycota), indicating that *Phyllotis* feeds on saxicolous lichen, as has been documented in other arctic and alpine mammals during periods of food scarcity (Conner, 1983; Seward, 2008; Richardson & Young, 1977). During the Arctic Winter, cricetid rodents such as snow voles (*Chionomys nivalis*) and northern bog lemmings (*Synaptomys borealis*) feed on tundra lichen (Richardson & Young, 1977). Likewise, during Winter months in the high alpine, North American pikas (*Ochotona princeps*) feed on lichens under the snowpack (Conner, 1983). Terricolous, arboreal, and saxicolous lichens are an important component of the winter diet of Caribou (*Rangifer tarandus*) in the northern Holarctic (Seaward, 2008) and arboreal lichens are an important component of the winter diet of Yunnan snub-nosed monkeys (*Rhinopithecus bieti*) in montane coniferous forests at elevations >4000 m (Kirkpatrick et al., 1998). Although lichens may serve as a seasonal or short-term supplement to the normal diet of many mammals living in arctic and alpine environments, their low nutritive value suggest that they are unlikely to represent a year-round dietary staple for small mammals like *Phyllotis* that have high metabolic demands.

We found no strong support for the arthropod fallout hypothesis, as we did not detect arthropod DNA in the gut contents of our summit mouse. In contrast to high-elevation pikas on the Quinghai-Tibetan Plateau that feed on yak feces, we found no evidence for interspecific coprophagy in our high-elevation *Phyllotis*, as indicated by the absence of metagenomic sequence reads and *COI* barcodes matching vicuña, guanaco, or any other potentially co-distributed mammals. The absence of such sequences also constitutes absence of evidence for scavenging.

### 3.4 Stable isotope analysis

Mice showed considerable variation in values of all three of the stable isotopes examined (Figure 5a-c, Table 2). Patterns included isotopic variation both among sites (indicating an elevational effect) and within sites (indicating individual variation in foraging habits). In the total dataset, δ^13^C values of *P. vaccarum* (Figure 5a, Table 2) varied between -22.8 and -12.7 ‰, with more ^13^C-enriched values being recorded at lower elevations (<4000 m). However, mice with relatively ^13^C-depleted values were captured at both sites 1 and 2. There was strong statistical support for inter-site differences in δ^13^C (PERMANOVA: PseudoF_5,34_ = 36.6, P_9999 perms_ = 0.0001). *Post-hoc* comparisons indicated significant differences (*P* < 0.05) between δ^13^C values for all sites apart from sites 3 and 4 (*t* = 1.42, *P* = 0.13), sites 3 and 6 (*t* = 1.92, *P* = 0.05), and sites 4 and 5 (*t* = 1.4, *P* = 0.14). *Phyllotis vaccarum* liver δ^15^N ranged between 6.2 and 22.8 ‰ (Figure 5b, Table 2), with considerable variation among sites (PseudoF_5,34_ = 39.3, P_9999 perms_ = 0.0001). Values were notably ^15^N-enriched at site 1 but included one individual with relatively low δ^15^N. *Post-hoc* comparisons showed overlap in δ^15^N values at sites 2 and 6 (*t* = 0.10, *P* = 0.92), sites 3 and 4 (*t* = 0.05, *P* = 0.97), sites 3 and 6 (*t* = 2.27, *P* = 0.05), and sites 4 and 5 (*t* = 0.97, *P* = 0.34). In the cases of both δ^13^C and δ^15^N, *P. vaccarum* showed a similar pattern of ^13^C- and ^15^N-enriched values at sites from lower elevations (< 4000 m) relative to individuals captured between 4000 and 5000 m (Figure 6, Table 2). This contrasted with the pattern in δ^34^S (Figure 5c, Table 2) where *P. vaccarum* showed less variation in general, with values ranging between -2.5 and 2.4 ‰, but exhibited a positive shift between ^34^S-depleted values at sites <4000 m and ^34^S-enriched values >4000 m. Values of δ^34^S for mouse livers varied among sites (PseudoF_5,34_ = 25.3, P_9999 perms_ = 0.0001), although variation was lower than that observed for C and N. *Post-hoc* comparisons indicated that δ^34^S values were similar for mice captured from sites 1 and 2 (*t* = 0.93, *P* = 0.70), sites 3 and 5 (*t* = 1.39, *P* = 0.18), sites 3 and 6 (*t* = 0.67, *P* = 0.52), sites 4 and 5 (*t* =1.35, *P* = 0.21), sites 4 and 6 (*t* = 1.8, *P* = 0.09, and sites 5 and 6 (*t* = 0.84, *P* = 0.45).

**Table 2.** Summary statistics (n, mean ± SD) for *Phyllotis vaccarum* liver stable isotope analyses of carbon (values shown for δ^13^C and lipid-corrected δ^13^C), nitrogen (δ^15^N), and sulfur (δ^34^S) and the elemental C:N ratio.

| Site | Elevation<br>(m) | n | $\delta^{13}\text{C}$<br>(‰) | Lipid corrected<br>$\delta^{13}\text{C}$ (‰) | $\delta^{15}\text{N}$<br>(‰) | $\delta^{34}\text{S}$<br>(‰) | C:N |
| --- | --- | --- | --- | --- | --- | --- | --- |
| Site 1 | 2370 | 8 | -18.3 (± 2.2) | -16.9 (± 2.0) | 19.2 (± 3.6) | -1.6 (± 0.8) | 4.2 (± 0.8) |
| Site 2 | 3240 | 8 | -16.2 (± 1.8) | -14.9 (± 1.8) | 10.1 (± 0.9) | -1.5 (± 0.7) | 4.0 (± 0.6) |
| Site 3 | 4150 | 4 | -22.6 (± 0.4) | -22 (± 0.2) | 7.6 (± 0.8) | 0.5 (± 1.3) | 3.4 (± 0.1) |
| Site 4 | 4360 | 8 | -22.8 (± 0.5) | -21.8 (± 0.2) | 7.6 (± 1.0) | 1.6 (± 0.5) | 3.8 (± 0.3) |
| Site 5 | 4620 | 5 | -23.3 (± 0.2) | -22.2 (± 0.4) | 7.1 (± 0.1) | 1.3 (± 0.1) | 3.7 (± 0.2) |
| Site 6 | 5070 | 7 | -22.0 (± 1.1) | -20.9 (± 0.9) | 10.2 (± 2.1) | 1.0 (± 0.9) | 3.8 (± 0.4) |
| Site 7 | 6739 | 1 | -22.0 (–) | -21.5 (–) | 7.0 (–) | 2.0 (–) | 3.3 (–) |

**Figure 5.**
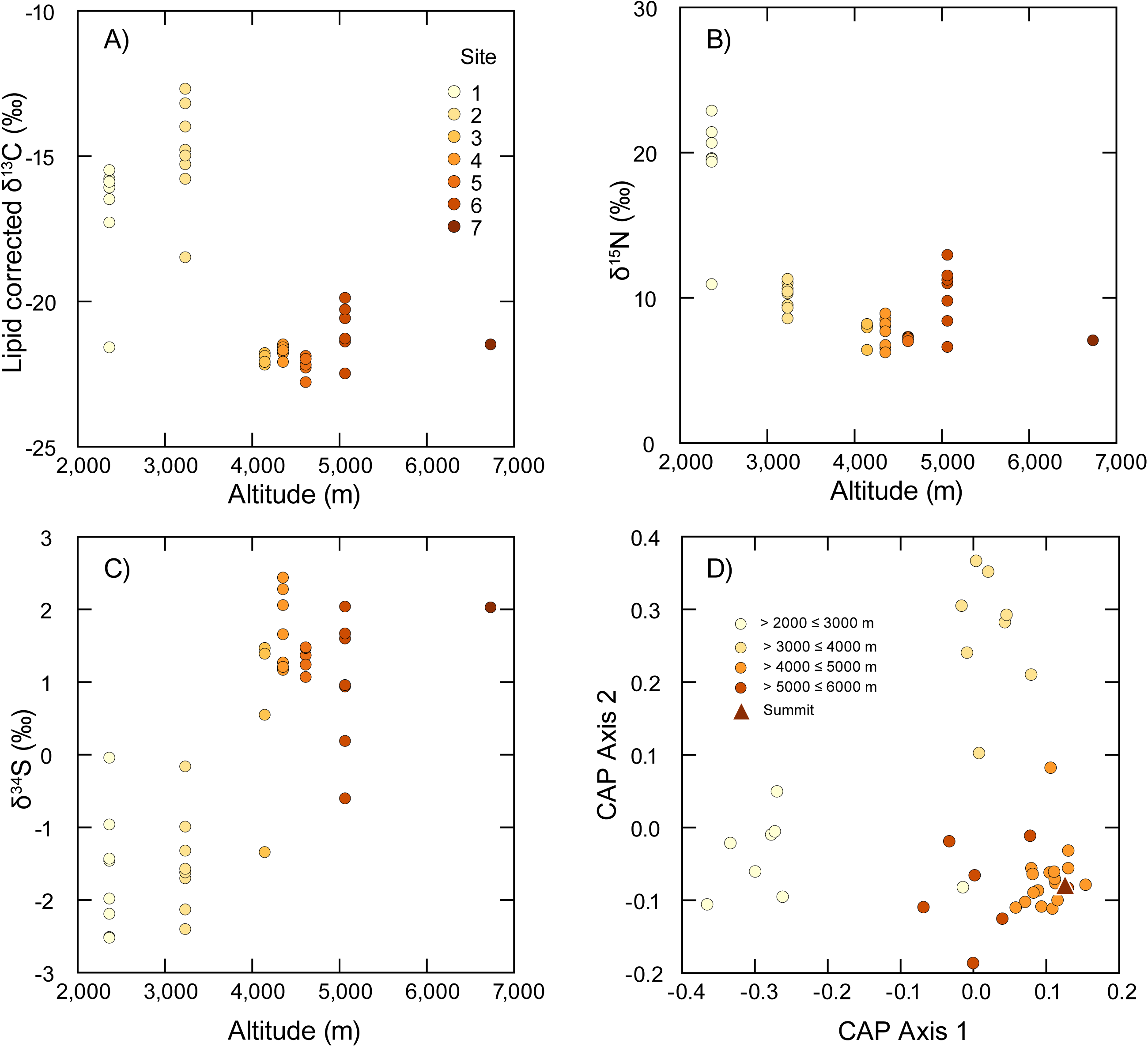
Variation in stable isotopes in livers of *Phyllotis vaccarum* sampled from different elevational zones: (A) δ^13^C, (B) δ^15^N, and (C) δ^34^S. (D) Results of multivariate CAP ordination based on Euclidean distances calculated from combined δ^13^C, δ^15^N and δ^34^S values.

**Figure 6.**
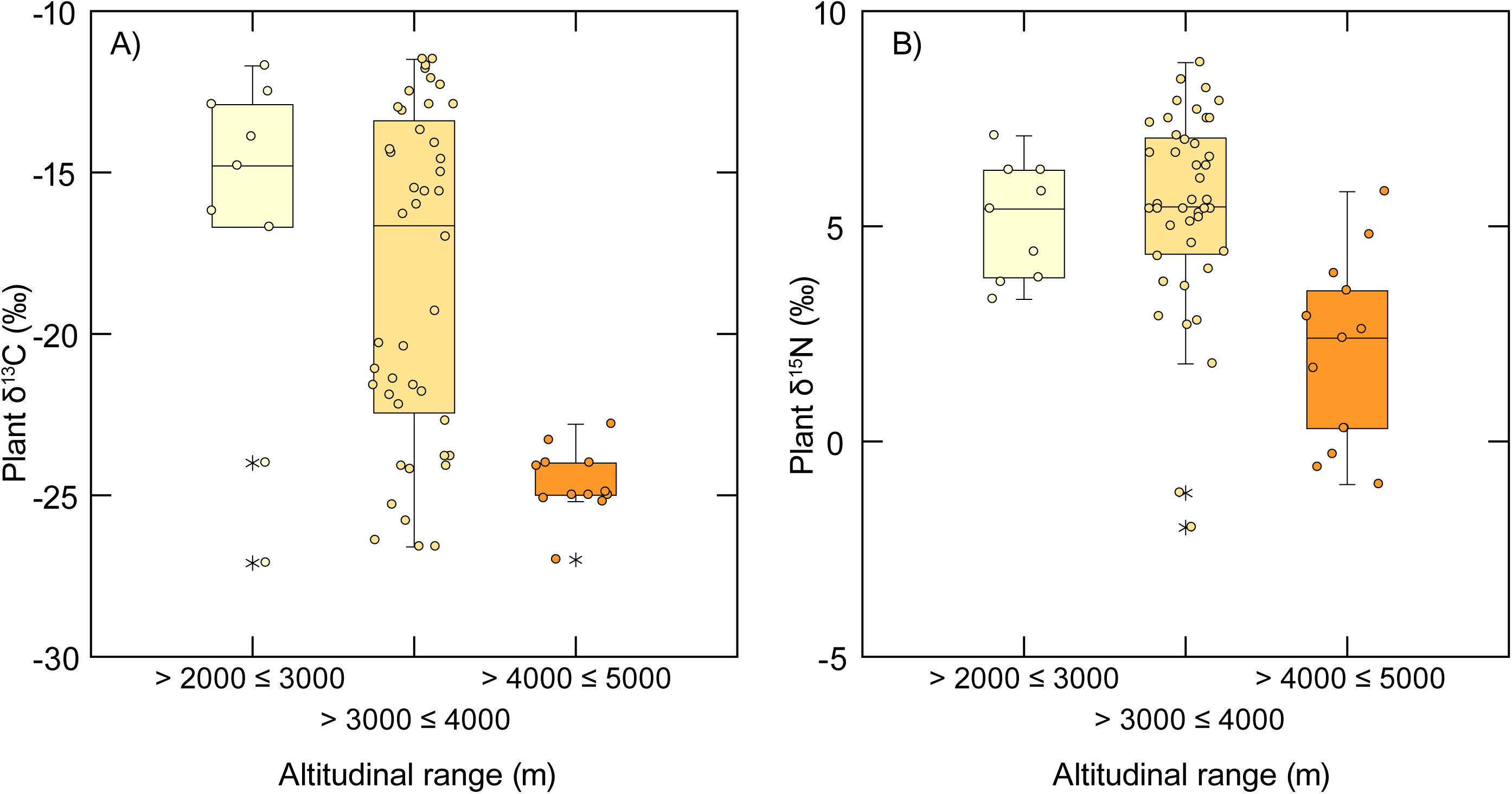
Variation in plant δ^13^C (A) and δ^15^N (B) across different elevational zones. Values taken from Díaz et al. (2016). Note the elevational shift in δ^13^C values showing dominance of C_4_ plants at lower elevations, a mix of C3, C4 and CAM plants at mid-elevations and a shift to C3 plants at higher elevations. Plant δ^15^N values were similar at lower and mid-elevations but were relatively ^15^N depleted at higher elevations. These data were used to define isotopic baselines for estimates of trophic position of *P*. *vaccarum*.

Mice from the highest elevations did not exhibit negative δ^15^N values, suggesting that lichenivory is not especially common.

#### 3.4.1 Assignment to capture elevations

As abiotic and biotic conditions (e.g. temperature, aridity, UV concentrations, plant nutrient availability, soil organic content) change with elevation, so too do the biomass and community composition of the primary producers (Díaz et al., 2019), with consequent changes in the availability of food for consumers and isotopic shifts at the base of the food web (Díaz *et al*. 2016). Given the known elevational gradient in stable isotope values, we can expect that stable isotope values will provide a means of identifying variation in habitat use among mice captured at different elevations.

Analysis of the combined stable isotope dataset using CAP showed that mice could be reliably assigned to broad elevational zones using individual δ^13^C, δ^15^N and δ^34^S values (Figure 5d) (CAP: Trace = 1.71, P = 0.0001). The leave-one-out classification cross validation (Table 3) indicated that by using the three isotope ratios we could assign an individual mouse to a 1000 m interval with ∼85 ‰ success. The CAP model predicted that the 6739 m summit mouse from Volcán Llullaillaco was isotopically most similar to mice captured from the 4000-5000 m interval (Figure 5d). The CAP model also suggested that 4 other individuals had stable isotope values characteristic of elevational zones distinct from where they were captured (Table 3). Including the summit mouse, this indicates that ca. 15 ‰ of the *P. vaccarum* in the study area had stable isotope values suggestive of upslope or downslope dispersal.

**Table 3.** Results of the canonical analysis of principal coordinates (CAP) leave-one-out cross-validation to assess the ability of the model to assign individual mice to the 1000 m elevational zone in which they were captured.

| Known capture altitude (m) | Predicted capture altitude (m) |  |  |  | % correctly classified |
| --- | --- | --- | --- | --- | --- |
|  | 2000-3000 | 3000-4000 | 4000-5000 | 5000-6000 |  |
| 2000 – 3000 | 7 | 0 | 0 | 1 | 86 |
| 3000 – 4000 | 0 | 7 | 0 | 1 | 86 |
| 4000 – 5000 | 0 | 0 | 16 | 1 | 94 |
| 5000 – 6000 | 0 | 0 | 2 | 5 | 71 |

#### 3.4.2 Trophic Position

Modal estimates of *P*. *vaccarum* trophic position (TP) varied between capture sites (Table 4) and ranged from 1.9 at site 2 to 4.3 at site 1. The latter estimate is extremely high and reflects the very high δ^15^N values from mice collected at site 1 (mean δ^15^N = 19.2 ‰). Discounting the results from site 1, mouse trophic position estimates were generally similar across sites and were indicative of omnivory with modal values between 1.9 and 2.3 at sites 2 to 5. The modal estimate for mice at site 6 was slightly higher (TP = 3.4), but the credibility limits overlapped with those from all sites apart from site 2. TP values between 2-3 are indicative of an herbivorous diet that includes some animal prey. TP values >3, as seen at site 6, indicates a diet dominated by animal prey. The TP of the summit mouse from Volcán Llullaillaco was estimated as 2.2, quite close to that of mice from sites 3, 4 and 5.

**Table 4.** Estimates of trophic position for mice captured in different elevational zones. Summary statistics are provided as modal TP (95 ‰ credibility intervals) and as mean TP ± SD.

| | *tRophicPosition*<br>model | Baseline data<br>(Díaz et al., 2016) | Trophic position<br>(mode, 95 % credibility<br>intervals) | Trophic<br>position<br>(mean ± SD) |
| --- | --- | --- | --- | --- |
| **Site 01** | *oneBaseline* | C3 & CAM data<br>combined from 2000<br>– 3000 m | <sup>†</sup>4.3 (3.4 – 5.1) | *4.3 ± 0.2 |
| **Site 02** | *twoBaselines* | C3 & CAM/C4 data<br>3000 – 4000 m | 1.9 (1.7 – 2.2) | 1.9 ± 0.1 |
| **Site 03** | *oneBaseline* | Plants >4000 m (all<br>C3) | 2.3 (1.8 – 2.9) | 2.3 ± 0.3 |
| **Site 04** | *oneBaseline* | Plants >4000 m (all<br>C3) | 2.3 (1.9 – 2.7) | 2.3 ± 0.2 |
| **Site 05** | *oneBaseline* | Plants >4000 m (all<br>C3) | 2.2 (1.8 – 2.5) | 2.2 ± 0.2 |
| **Site 06** | *oneBaseline* | Plants >4000 m (all<br>C3) | 2.9 (2.3 – 3.5) | 2.9 ± 0.3 |
| **Summit**<br>**mouse** | *oneBaseline-greta* | Plants >4000 m (all<br>C3) | 2.2 (1.9 – 2.4) | 2.2 ± 0.1 |

<sup>†</sup> $\delta^{15}\text{N}$ values of mice captured at Site 1 were unusually high and were markedly $^{15}\text{N}$ enriched relative to the putative baseline (see Figure 5b) and may be artefactual.

## 4. CONCLUSIONS

A combination of metagenomic, metabarcoding, and stable isotope data provided new insights into the diet of *Phyllotis* mice living at extreme elevations that far surpass known vegetation limits. Stable isotope data revealed that *Phyllotis vaccarum* maintains a mainly omnivorous diet in all elevational zones, and elevational variation in diet reflects variation in vegetation composition and the extent to which the mice rely on animal prey. Estimates of trophic position based on isotopic data indicated that mice collected near apparent vegetation limits (∼5100 m) on the flanks of Llullaillaco rely more heavily on animal prey than mice from lower elevations. Metagenomic and metabarcoding analyses of gut contents from the mouse from the summit of Llullaillaco (6739 m) revealed a strictly herbivorous diet. The absence of animal DNA suggests that mice at extreme elevations do not subsist on wind-blown arthropods or other animal material. The detection of DNA from lichen-associated fungi indicate that *Phyllotis* mice living above known vegetation limits may supplement their diet with saxicolous lichens, as observed for other arctic and alpine mammals during periods of food scarcity in the winter. However, measured δ^15^N levels indicate that lichen is not an important dietary staple in mice native to any of the surveyed elevational zones. The metagenomic and metabarcoding data also produce a scientific conundrum: the gut contents of a mouse captured at 6739 m elevation contained DNA from several families of native plants that are not known to occur above ∼5000 m elevation. Clearly we have more to learn about the elevational distributions of both the plants and the mice. It is possible that some of the plants identified in the diet of the summit mouse exist at higher elevations than previously supposed, but they exist beneath the snowpack or in other cryptic microhabitats.

## AUTHOR CONTRIBUTIONS

CQ-R, CH, and JFS designed the study, MQC, GD, and JFS performed the fieldwork, CQ-R and CH performed data analysis, CQ-R, CH, and JFS wrote the initial draft of the manuscript, and all authors read and approved it.

## ACKNOWLEDGMENTS

We thank Mario Pérez-Mamani and Juan Carlos Briceño for assistance and companionship in the field, José Urquizo for help with figures, and Cristina Dorador for logistical assistance.

