## Supplemental Information for "Diet of Andean leaf-eared mice (*Phyllotis*) living at extreme elevations on Atacama volcanoes: insights from metagenomics, DNA metabarcoding, and stable isotopes"

Tables S1-S6

**Table S1**. Input, filtered, denoised forward, denoised reverse, merged and non-chimeric reads for trnL P6 loop primers per sample.

| Sample | Input | Filtered | DenoisedF | DenoisedR | Merged | Nonchim |
| --- | --- | --- | --- | --- | --- | --- |
| Extraction blank | 17 | 10 | 9 | 7 | 7 | 7 |
| stomach | 65.723 | 63.552 | 63.537 | 63.504 | 61.611 | 61.491 |
| cecum | 85.849 | 71.919 | 71.819 | 71.622 | 69.401 | 68.688 |
| C1 | 87.333 | 82.656 | 82.563 | 82.579 | 80.032 | 80.032 |
| C2 | 66.839 | 64.199 | 64.185 | 64.186 | 62.649 | 62.649 |
| C3 | 70.175 | 67.173 | 66.928 | 67.042 | 64.984 | 64.984 |
| C4 | 69.287 | 66.445 | 66.367 | 66.427 | 64.282 | 64.282 |
| C5 | 64.639 | 62.033 | 61.992 | 61.949 | 59.975 | 59.836 |
| C6 | 72.401 | 69.428 | 69.406 | 69.400 | 67.528 | 67.528 |
| C7 | 77.599 | 73.226 | 73.059 | 73.050 | 70.801 | 70.797 |
| C8 | 71.317 | 64.527 | 64.444 | 64.344 | 62.261 | 62.261 |
| C9 | 75.808 | 72.965 | 72.928 | 72.883 | 71.345 | 70.517 |
| C10 | 66.679 | 59.690 | 59.569 | 59.357 | 57.510 | 57.475 |
| C11 | 59.982 | 56.208 | 56.078 | 56.082 | 54.377 | 54.377 |
| C12 | 66.209 | 62.673 | 62.581 | 62.557 | 60.312 | 60.312 |
| **TOTAL** | **999.857** | **936.704** | **935.465** | **934.989** | **907.075** | **905.236** |

**Table S2**. Input, filtered, denoised forward, denoised reverse, merged and non-chimeric reads for ITS2 primers per sample.

| Sample | Input | Filtered | DenoisedF | DenoisedR | Merged | Nonchim |
| --- | --- | --- | --- | --- | --- | --- |
| stomach | 186.662 | 51.882 | 51.729 | 51.665 | 43.165 | 40.150 |
| cecum | 192.646 | 38.775 | 38.675 | 38.576 | 28.196 | 27.172 |
| C1 | 52.978 | 12.036 | 11.994 | 11.989 | 8.971 | 8.969 |
| C2 | 131.565 | 23.473 | 23.366 | 23.276 | 18.090 | 18.049 |
| C3 | 96.694 | 21.695 | 21.650 | 21.587 | 16.442 | 16.436 |
| C4 | 123.788 | 24.462 | 24.379 | 24.344 | 18.907 | 18.713 |
| C5 | 69.594 | 15.980 | 15.914 | 15.912 | 11.709 | 11.289 |
| C6 | 104.771 | 23.918 | 23.846 | 23.812 | 18.447 | 18.441 |
| C7 | 126.643 | 32.258 | 32.161 | 32.020 | 24.754 | 24.652 |
| C8 | 122.089 | 32.204 | 32.076 | 31.980 | 25.172 | 24.829 |
| C9 | 95.100 | 31.228 | 31.127 | 31.081 | 25.353 | 25.026 |
| C10 | 92.587 | 22.065 | 21.927 | 21.975 | 16.832 | 16.639 |
| C11 | 85.107 | 22.338 | 22.236 | 22.172 | 17.711 | 17.570 |
| C12 | 69.403 | 18.381 | 18.328 | 18.274 | 14.278 | 14.036 |
| **TOTAL** | **1.549.627** | **370.695** | **369.408** | **368.663** | **288.027** | **281.971** |

**Table S3**. Input, filtered, denoised forward, denoised reverse, merged and non-chimeric reads for p23SrV primers per sample.

| Sample | input | filtered | denoisedF | denoisedR | merged | nonchim |
| --- | --- | --- | --- | --- | --- | --- |
| Extraction blank | 116 | 18 | 15 | 15 | 0 | 0 |
| stomach | 164.960 | 34.357 | 34.303 | 34.285 | 18.684 | 18.608 |
| cecum | 159.183 | 29.170 | 28.900 | 28.876 | 24.067 | 23.714 |
| C1 | 105.590 | 18.439 | 18.325 | 18.292 | 14.736 | 14.717 |
| C2 | 108.051 | 10.336 | 10.249 | 10.232 | 8.665 | 8.662 |
| C3 | 144.892 | 34.201 | 34.058 | 33.990 | 26.655 | 26.598 |
| C4 | 91.943 | 15.960 | 15.854 | 15.791 | 12.904 | 12.859 |
| C5 | 126.028 | 18.813 | 18.679 | 18.668 | 15.074 | 14.965 |
| C6 | 153.258 | 28.918 | 28.684 | 28.567 | 23.668 | 23.513 |
| C7 | 132.896 | 19.196 | 18.973 | 18.880 | 15.059 | 14.903 |
| C8 | 158.723 | 30.256 | 29.972 | 29.933 | 24.503 | 24.099 |
| C9 | 168.609 | 31.725 | 31.394 | 31.382 | 25.961 | 24.907 |
| C10 | 140.728 | 19.920 | 19.727 | 19.650 | 16.087 | 15.898 |
| C11 | 119.463 | 24.898 | 24.625 | 24.547 | 19.867 | 19.658 |
| C12 | 160.913 | 38.832 | 38.473 | 38.418 | 33.246 | 32.837 |
| **TOTAL** | **1.935.353** | **355.039** | **352.231** | **351.526** | **279.176** | **275.938** |

**Table S4**. Input, filtered, denoised forward, denoised reverse, merged and non-chimeric reads for ITS1 primers per sample.

| Sample | input | filtered | denoisedF | denoisedR | merged | nonchim |
| --- | --- | --- | --- | --- | --- | --- |
| Extraction blank | 30 | 14 | 5 | 3 | 3 | 3 |
| stomach | 85.919 | 60.669 | 60.285 | 60.359 | 58.840 | 57.953 |
| cecum | 85.667 | 55.148 | 54.810 | 54.640 | 53.480 | 53.017 |
| C1 | 100.641 | 57.430 | 57.176 | 57.123 | 54.667 | 50.079 |
| C2 | 149.940 | 97.006 | 96.686 | 96.554 | 90.605 | 85.856 |
| C3 | 95.950 | 61.266 | 61.019 | 60.872 | 58.095 | 57.108 |
| C4 | 88.134 | 57.532 | 57.335 | 57.250 | 54.658 | 53.568 |
| C5 | 64.992 | 37.085 | 36.865 | 36.754 | 35.340 | 35.011 |
| C6 | 69.510 | 46.261 | 45.984 | 45.761 | 41.148 | 38.846 |
| C7 | 93.476 | 52.639 | 52.279 | 52.107 | 49.869 | 45.908 |
| C8 | 119.736 | 88.616 | 88.288 | 88.184 | 86.796 | 84.821 |
| C9 | 120.747 | 78.783 | 78.313 | 78.173 | 75.869 | 74.128 |
| C10 | 98.785 | 63.102 | 62.659 | 62.609 | 60.707 | 59.641 |
| C11 | 141.305 | 74.558 | 74.107 | 73.899 | 64.854 | 62.434 |
| C12 | 183.546 | 100.844 | 100.256 | 100.094 | 97.551 | 95.722 |
| **TOTAL** | **1.498.378** | **930.953** | **926.067** | **924.382** | **882.482** | **854.095** |

**Table S5**. Input, filtered, denoised forward, denoised reverse, merged and non-chimeric reads for COI (metazoan) primers per sample.

| Sample | input | filtered | denoisedF | denoisedR | merged | nonchim |
| --- | --- | --- | --- | --- | --- | --- |
| Extraction blank | 1.816 | 1.117 | 1.116 | 1.114 | 0 | 0 |
| stomach | 166.575 | 79.737 | 79.626 | 79.668 | 1.413 | 1.412 |
| cecum | 133.126 | 71.365 | 71.264 | 71.288 | 840 | 819 |
| C1 | 101.655 | 55.856 | 55.789 | 55.840 | 874 | 873 |
| C2 | 138.917 | 78.527 | 78.461 | 78.501 | 669 | 669 |
| C3 | 130.863 | 79.014 | 78.922 | 78.954 | 2.610 | 2.610 |
| C4 | 90.177 | 32.118 | 32.076 | 32.085 | 796 | 796 |
| C5 | 169.215 | 101.419 | 101.294 | 101.365 | 2.129 | 2.129 |
| C6 | 155.282 | 86.844 | 86.756 | 86.758 | 1.685 | 1.685 |
| C7 | 146.412 | 83.350 | 83.184 | 83.233 | 3.704 | 3.604 |
| C8 | 169.047 | 92.995 | 92.885 | 92.918 | 1.821 | 1.811 |
| C9 | 150.635 | 73.805 | 73.715 | 73.746 | 725 | 725 |
| C10 | 161.401 | 90.767 | 90.645 | 90.656 | 3.411 | 3.411 |
| C11 | 155.326 | 88.127 | 87.905 | 87.800 | 3.250 | 3.230 |
| C12 | 135.045 | 69.068 | 69.003 | 69.010 | 1.802 | 1.802 |
| **TOTAL** | **2.005.492** | **1.084.109** | **1.082.641** | **1.082.936** | **25.729** | **25.576** |

**Table S6**. Input, filtered, denoised forward, denoised reverse, merged and non-chimeric reads for COI (invertebrates) primers per sample.

| Sample | input | filtered | denoisedF | denoisedR | merged | nonchim |
| --- | --- | --- | --- | --- | --- | --- |
| Extraction blank | 30 | 14 | 5 | 3 | 3 | 3 |
| stomach | 85.919 | 60.669 | 60.285 | 60.359 | 58.840 | 57.953 |
| cecum | 85.667 | 55.148 | 54.810 | 54.640 | 53.480 | 53.017 |
| C1 | 100.641 | 57.430 | 57.176 | 57.123 | 54.667 | 50.079 |
| C2 | 149.940 | 97.006 | 96.686 | 96.554 | 90.605 | 85.856 |
| C3 | 95.950 | 61.266 | 61.019 | 60.872 | 58.095 | 57.108 |
| C4 | 88.134 | 57.532 | 57.335 | 57.250 | 54.658 | 53.568 |
| C5 | 64.992 | 37.085 | 36.865 | 36.754 | 35.340 | 35.011 |
| C6 | 69.510 | 46.261 | 45.984 | 45.761 | 41.148 | 38.846 |
| C7 | 93.476 | 52.639 | 52.279 | 52.107 | 49.869 | 45.908 |
| C8 | 119.736 | 88.616 | 88.288 | 88.184 | 86.796 | 84.821 |
| C9 | 120.747 | 78.783 | 78.313 | 78.173 | 75.869 | 74.128 |
| C10 | 98.785 | 63.102 | 62.659 | 62.609 | 60.707 | 59.641 |
| C11 | 141.305 | 74.558 | 74.107 | 73.899 | 64.854 | 62.434 |
| C12 | 183.546 | 100.844 | 100.256 | 100.094 | 97.551 | 95.722 |
| **TOTAL** | **1.498.378** | **930.953** | **926.067** | **924.382** | **882.482** | **854.095** |
